# Parliament2: Fast Structural Variant Calling Using Optimized Combinations of Callers

**DOI:** 10.1101/424267

**Authors:** Samantha Zarate, Andrew Carroll, Olga Krashenina, Fritz J Sedlazeck, Goo Jun, William Salerno, Eric Boerwinkle, Richard Gibbs

## Abstract

Here we present Parliament2 – a structural variant caller which combines multiple best-in-class structural variant callers to create a highly accurate callset. This captures more events than the individual callers achieve independently. Parliament2 uses a call-overlap-genotype approach that is highly extensible to new methods and presents users the choice to run some or all of Breakdancer, Breakseq, CNVnator, Delly, Lumpy, and Manta to run. Parliament2 applies an additional parallelization framework to speed certain callers and executes these in parallel, taking advantage of the different resource requirements to complete structural variant calling much faster than running the programs individually. Parliament2 is available as a Docker container, which pre-installs all required dependencies. This allows users to run any caller with easy installation and execution. This Docker container can easily be deployed in cloud or local environments and is available as an app on DNAnexus.

## Introduction

Structural variants (SVs) are large (50bp+) variations of the genome [1,2]. Because these variations are similar to or most often larger than the size of individual short reads, they are difficult to detect directly in short-read data and must instead be inferred through several types of indirect signals, including split-read mapping, soft-clipping, changes in the distance between and orientation of read pairs, changes in coverage depth, or alterations in the heterozygosity of a region[3]. While accuracy of SNP and Indel calling pipelines can exceed 99.9%[4], accuracy of structural variant pipelines is reported as significantly lower, with the Genome in a Bottle[5] only recently generating benchmark sets.

Various structural callers, including Breakdancer [6], CNVnator [7], Crest [8], Delly [9] Lumpy [10], Manta [11], and Pindel [12] rely on different heuristics leveraging some or most of the mapped read signals to infer structural variants. This diversity of approaches causes certain programs to perform better or worse on structural variants of different types – e.g. Deletion (DEL), Insertion (INS), Duplication (DUP), Inversion (INV), and Translocation (TRA) – and across different size ranges.

This diversity in approaches and domain performance makes ensemble strategies appealing. Two methods: MetaSV [13] and Parliament1 [14], employ a three step Overlap-Merge-Validate strategy to combine results of multiple callers into a high-quality consensus set. Both MetaSV and Parliament1 use an assembly-based method for the validation step, which is very accurate in Parliament1 but is computationally intensive and limits the maximum size of events. Because MetaSV and Parliament1 start from already-generated structural variant calls, they place the burden of installing and running individual SV callers on the user.

In this paper, we present Parliament2, a refinement of Parliament1 designed to economically scale SV calling to 10,000 WGS cohort scale at tunable sensitivity and specificity. Parliament2 allows a user to run some, or all, of Breakdancer, Breakseq, CNVnator, Delly, Lumpy, and Manta to generate calls; uses SURVIVOR [15] to overlap these calls into consensus candidates; and validates these calls using SVTyper [16].

Parliament2 uses different parallelization strategies to run callers simultaneously, taking advantage of the different requirements in CPU, disk I/O, and RAM that each program has at a given time. This allows Parliament2 to complete all callers in roughly the same wall-clock and CPU time as the slowest individual caller, CNVnator. A 16-core machine can process a 35X Whole Genome Sequence (WGS) sample in 2-5 hours. Parliament2 produces both raw results for each individual caller, and a genotyped consensus VCF of all supported calls.

Parliament2 is open-source and available both as code base (https://github.com/dnanexus/parliament2) and as a Docker image which can be used to easily run any single, multiple, or all individual caller from Parliament2, (https://hub.docker.com/r/dnanexus/parliament2) Parliament2 is also a publicly available app on DNAnexus. This also allows a user to run only individual components if desired. Furthermore, Parliament2 is easily extendable allowing a user wanting to run a tool to tap into Parliament2’s accelerated execution. Optionally, Parliament2 provides PDF images of SVs and their supportive reads to enable a manual curation using SVVIZ [17].

## Results

### Parliament2 speeds execution with multiple parallelization strategies

Parliament2 uses two different parallelization approaches to run multiple callers at the greatest efficiency. Callers used by Parliament2 fall into three classes: those capable of parallelization by native multi-threading within the tool (Breakseq, Manta); those with built-in methods for parallelization by chromosome (CNVnator, Breakdancer); and those lacking either (Delly, Lumpy). Upon execution, Parliament2 immediately begins running Breakseq and Manta with multiple threads. It then splits the input BAM by chromosome and initiates runs on the remaining callers. This leads to more than a 10-fold reduction in runtime on a 35X WGS sample (Table 1) on a 16-core machine.

**Table 1.** Runtime speedup due to parallelization.

| Tool | Native Runtime | Parallelized Runtime |
| --- | --- | --- |
| LumpyExpress | 6.45 hours | 0.45 hours |
| Delly | 8.52 hours | 0.67 hours |

Parliament2 also parallelizes by running multiple programs concurrently. This leads to a reduction in runtime by achieving higher overall machine utilization of resources (Figure 1). In cluster and cloud environments, where machines are provisioned to conduct a specific task, this generally translates fully to saved compute resources and costs in addition to the saved wall-clock time.

**Figure 1.**
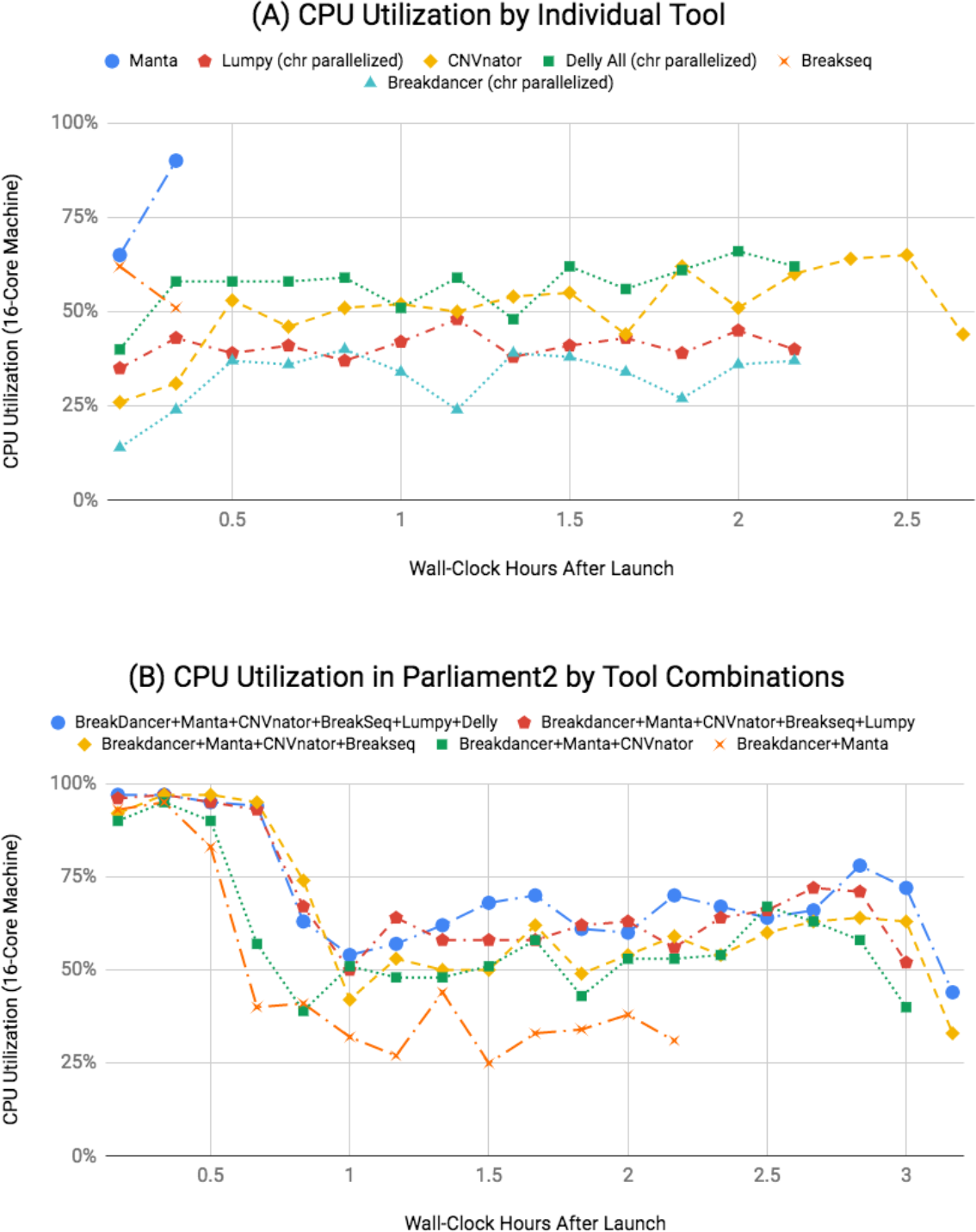
Concurrent execution of multiple tools in Parliament2 increases resource utilization. (A) Percent of total CPU utilization on a 16-core machine executing Parliament2 and running only an individual tool. Each line concludes when the program finishes executing. For example, Delly and Breakdancer each require about 2 hours to finish. (B) Resource utilization when running combinations of methods simultaneously within Parliament2. Starting from Breakdancer, Manta, and CNVnator (the green line), Breakseq, Lumpy, and Delly can be added (representing the blue line of all 6 callers) with only 10 minutes of additional wall-clock time.

### Parliament2 enables high-precision calls

To assess accuracy, we ran Parliament2 on 35X WGS of HG002 from the PrecisionFDA Truth Challenge. We compared these calls against the Genome in a Bottle v0.6 SV candidate truth set [18] using the Truvari comparator [19] for SV calls above 50bp. Parliament2 finished the analysis in 3 hours 26 minutes on a 16-core machine.

Parliament2 calls several different types of events: deletions, insertions, duplications, inversions, and translocations are all called, which most being called by several different types of callers. However, we focus most of our analysis on deletion events. We do so because the Genome in a Bottle truth set only contains validated calls of the deletion and insertion type. Also as we show later, discovery power with short-reads is much weaker at detecting insertions by any method.

When Parliament2 combines outputs from multiple callers, it records which callers support each call event. This allows calculation of the precision of calls made by any of the individual tools, or by combinations. The precision of individual methods ranges from 8% with CNVnator to 92% for Manta, the most precise caller. However, when a call is made by multiple methods, precision can reach 100% (Figure 2A).

Fewer of the individual methods call insertion events (Breakseq, Manta, and Delly). The precision of these calls is high (Figure 2B), with the intersection of Delly and Manta reaching 100% precision.

**Figure 2.**
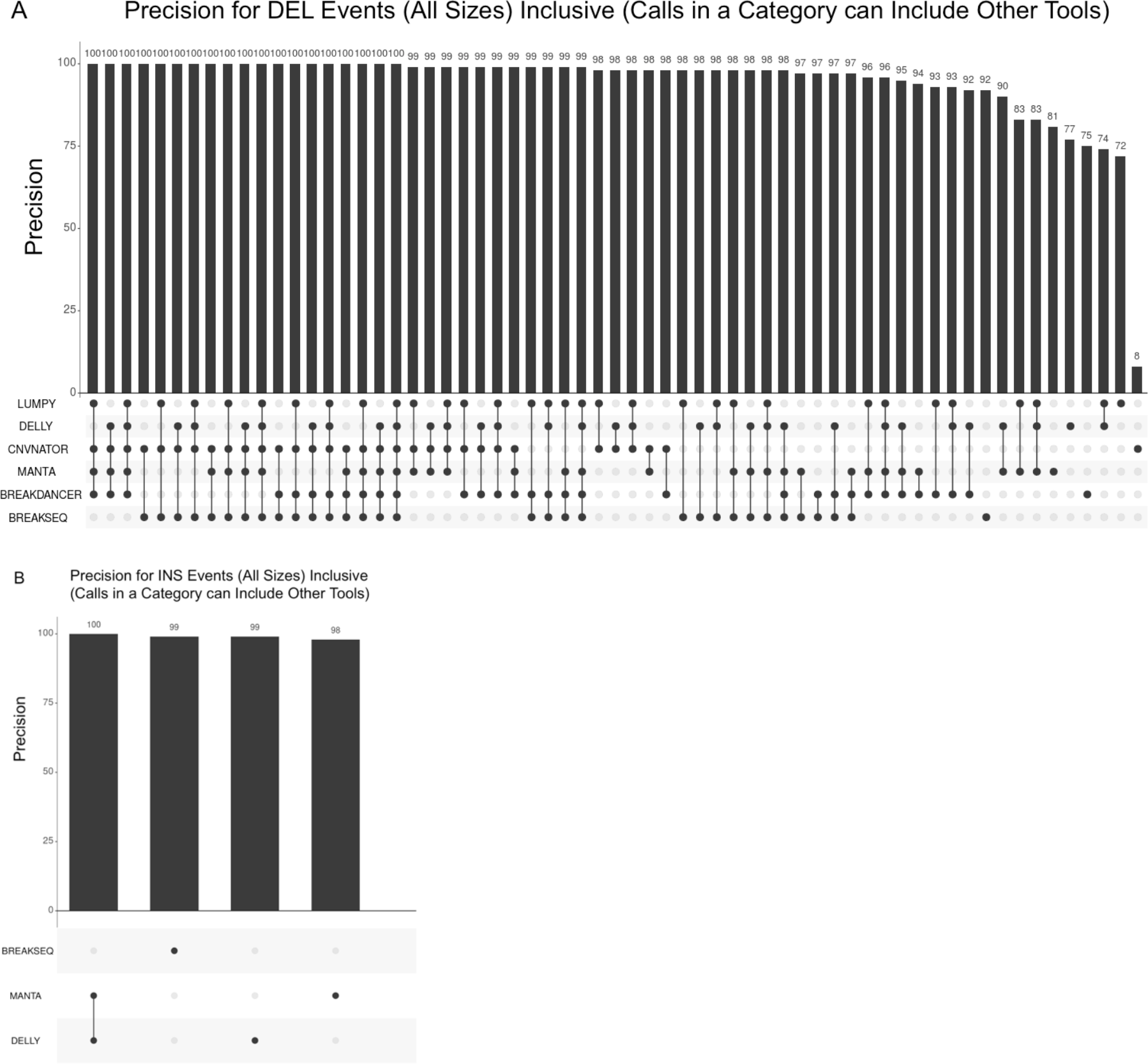
Precision of individual methods and combinations of methods. This Upset [20] plot shows the precision of calls supported by individual callers and combinations of callers for deletions (A) and insertions (B). Calls found in one caller and in the intersection of multiple callers are still included in the individual caller, allowing for a direct comparison of precision between the individual methods (i.e. calls supported by MANTA have, in aggregate, 92% precision regardless of other supporting callers).

### Comparison of caller overlaps with the Genome in a Bottle set allows filtering based on confidence

In the prior section, we expressed precision stratified across callers in an inclusive manner (i.e. any call present in a superset of one caller category was still included in precision numbers of the subset) (Figure 2A). This allows a comparison of the overall precision of individual methods such as Lumpy versus Delly. However, expressing the same data as exclusive (precision of calls which appear in Delly alone, not in either Delly and Lumpy) allows us to quantify the greater resolution of precision possible. For example, the overall precision of all Delly deletion calls is 77%, but the precision of calls found by Delly but not by any other method is 25%, allowing a substantially more granular estimate of call quality.

Figure 3 shows Parliament deletion calls and the combination of the exact supporting methods and their precision as calculated from the Genome in a Bottle v0.6 SV truth set. Based on this analysis, it is possible to filter events at below 50% precision based on which callers support the event. Parliament2 uses the LowQual filter to report events below this value.

**Figure 3.**
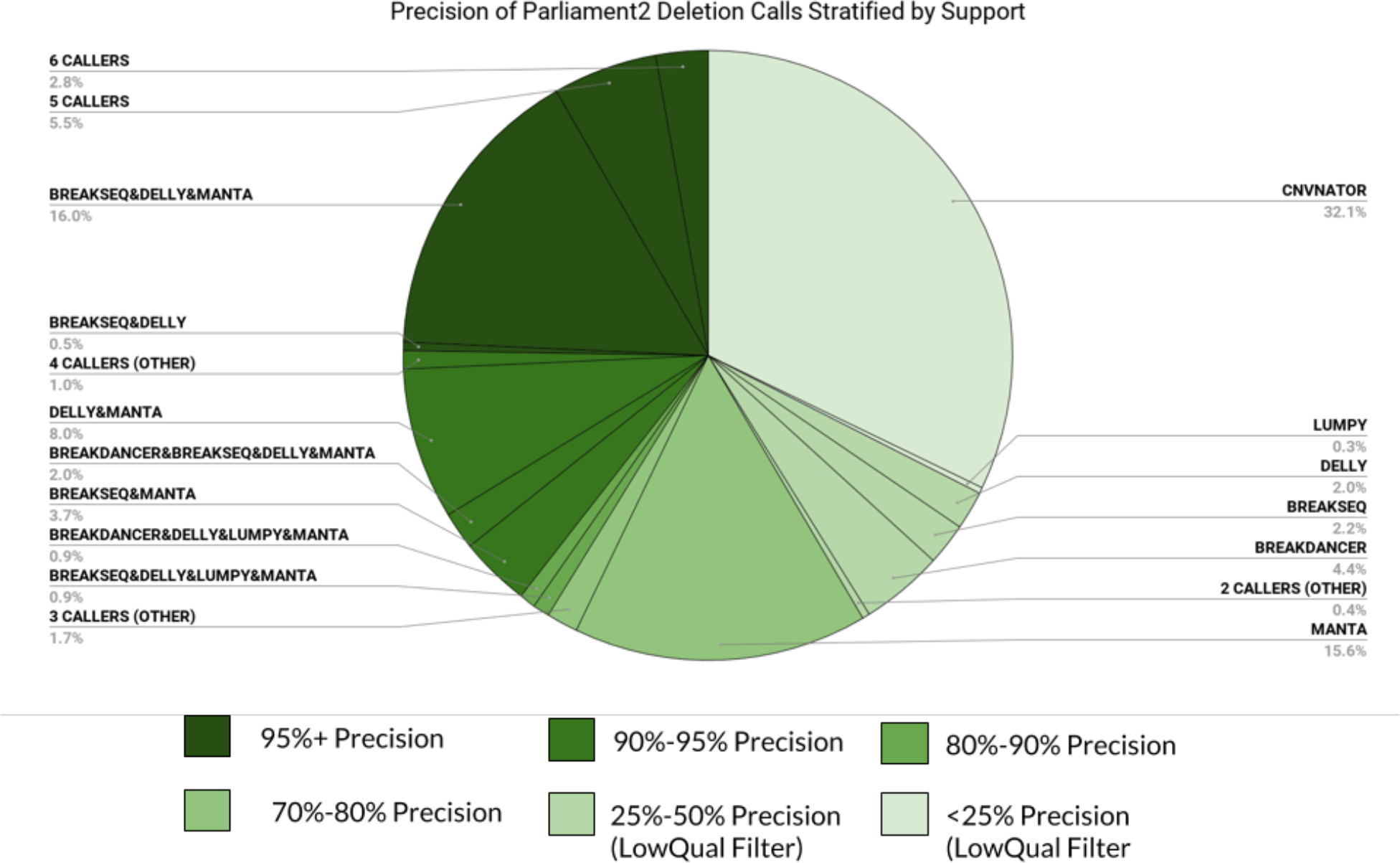
Precision of Parliament2 for DEL events stratified by support from specific callers. The figure above stratifies deletion calls produced by Parliament2 on a 35X HG002 WGS by the callers that support each event. The precision of each tier is labelled based on the precision calculated from the v0.6 Genome in a Bottle SV Truth Set. Combinations of callers with precision below 50% are reported as “LowQual” filter in Parliament2 output. Categories will fewer than 20 calls are combined together based on the number of supporting callers. The grey percentages indicate the fraction of total Parliament2 calls that fall into each category (e.g. 3.7 % of all Parliament2 calls are called by exactly Breakseq and Manta).

### Comparison of call overlaps allows assignment of quality values to each call

Parliament2 expresses the call quality as a Phred-encoded value within its final output VCF. This is based on the precision results from Genome in a Bottle for the exact combinations of supporting callers, the type of the event (deletion or insertion), and the size category of the event (50bp – 300bp, 300bp – 1kb, 1kb+). This quality value allows investigators to set thresholds to achieve the trade-off between precision and recall that is desired for their use case or to prioritize events based on how likely they are to be true events. Figure 4 shows the quality based on caller support and the size.

**Figure 4.**
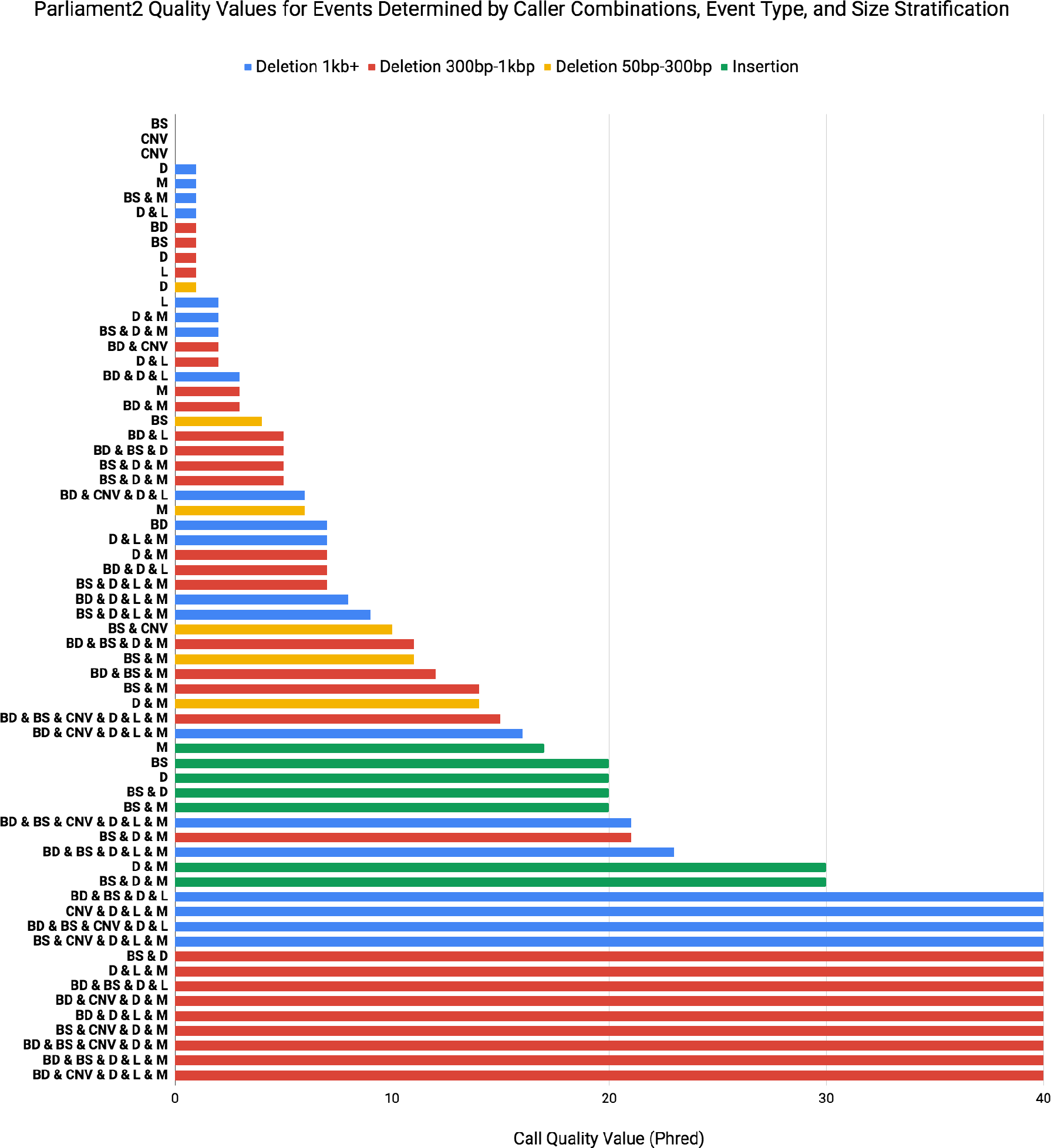
Quality values assigned by Parliament2 to SV events of various types, sizes, and support. Parliament2 assigns a quality value to each event based on the precision observed in comparisons with the Genome in a Bottle v0.6 truth set. In the above figure, the event type (deletion or insertion) and size determine the color code. The maximum QV assigned is 40, even if the precision of the subset is higher. Only categories with more than 2 calls are included.

### Parliament 2 increases recall by leveraging the specialization of multiple methods

Running multiple callers generally results in discovery of more candidate calls, in aggregate. In addition, different methods have different success in discovering events of different sizes. For example, Manta has the best recall for events from 50bp – 300 bp at 55% compared with Delly’s second highest recall at 27% (Figure 5A). However, for events larger than 1kb in size, Manta has only the fourth highest recall at 87% (Figure 5B), compared with the leader Delly and Lumpy at 93% and 92% respectively. Strong performance across multiple size ranges is important because while the larger number of small events tends to make precision and recall in the 50bp – 300 bp range dominate aggregate measures of accuracy, large events (1kpb+), though rarer, can have more significant biological impacts.

**Figure 5.**
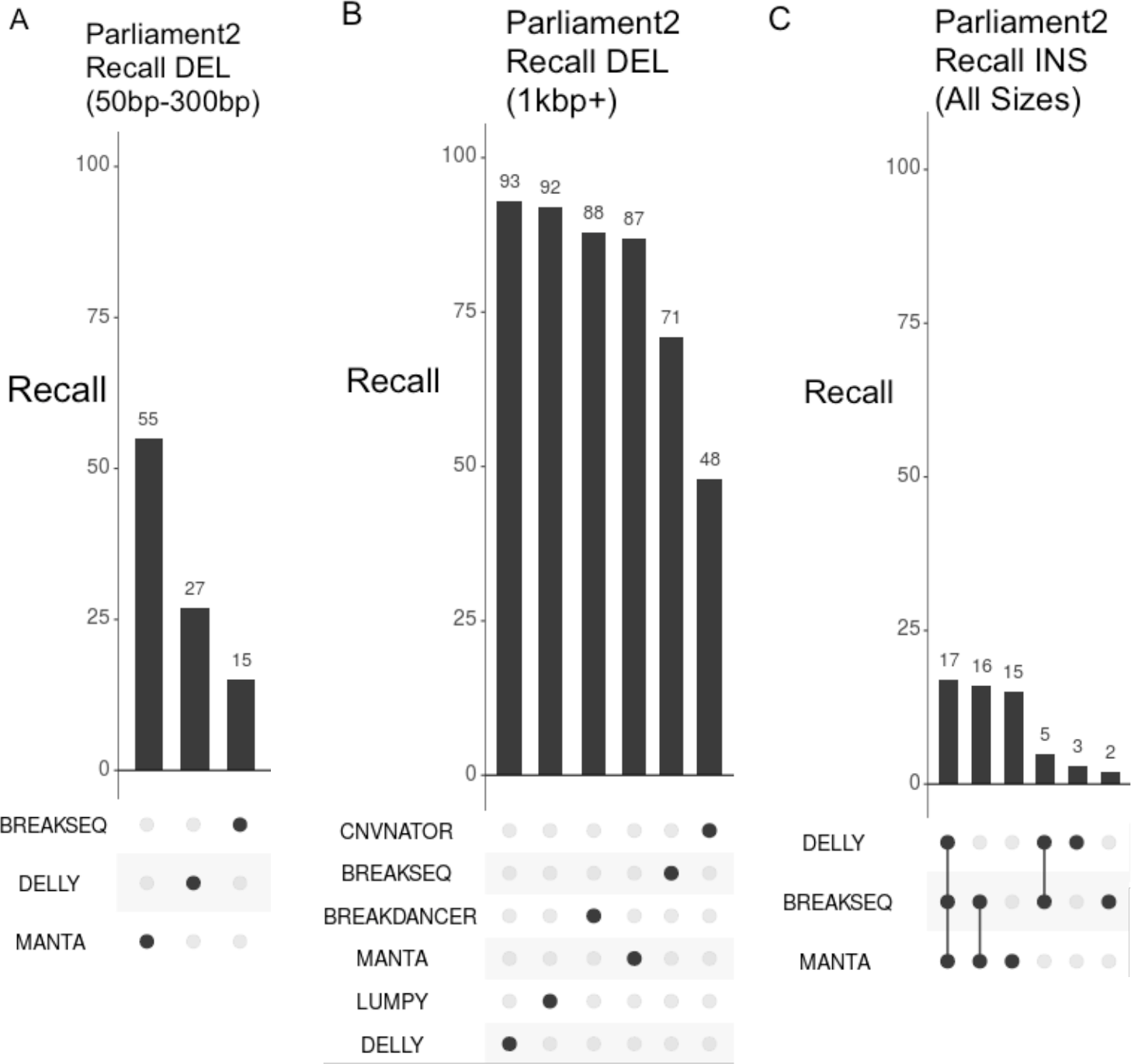
Recall of deletion events for on small and large events, and for insertions. Individual methods differ in their discovery power between (A) smaller deletion events and (B) larger deletions. (C) Recall for insertion events of all sizes is lower. The difference in discovery power by event type and size allows Parliament2 to discover more events in total

Discovery power for indels with these short-read methods is worse than for deletion events. Manta is by far the strongest method for detecting insertions, with Delly and Breakseq each increasing discovery power by 6.7% relative to Manta. Taking the combination of individual methods increases the overall recall that is possible for deletion events (Figure 6), reaching a total recall of 71%. Achieving a high recall is especially important for large population projects where it is possible to generate a discovery set of all individual samples as well as to return to individual samples to generate a force-call for every candidate discovered. This allows more refined methods for filtering after call generation (like looking at breakpoint consistency or Hardy-Weinberg equilibrium [21]).

**Figure 6.**
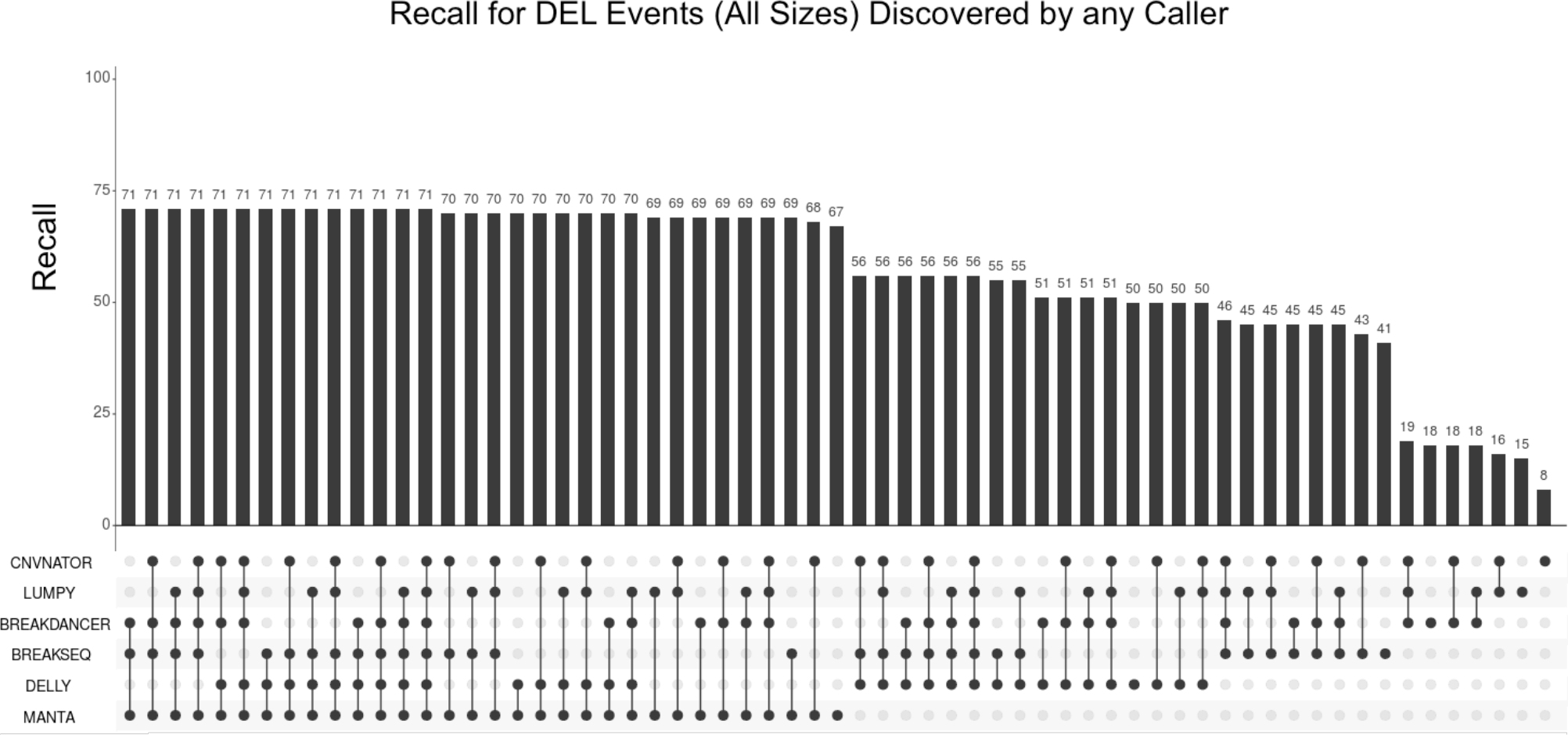
Recall of Parliament2 for DEL Events for an Event Found in any Sample. An Upset plot shows the recall achieved in the union of the callers in Parliament. The noticeable step-like increase marks the addition of a new individual caller. This may reflect the use of different discovery strategies by various methods giving access to certain events that can only be discovered through one type of signal (e.g. split-read, coverage, or insert size).

### Parliament2 Produces More Accurate Final Calls

The innovations discussed here: the ability to generate and combine multiple tools with diverse strengths, to use a known benchmark to develop quality stratification based on the specific support for events, and to filter events which fall below a threshold allow Parliament2 to create a final set of results which is greater than the sum of its parts. The thoughtful union of these multiple methods allows Parliament2 to discover more events across a variety of size ranges without a loss of precision by the inclusion of less precise methods (Figure 7). Note that the accuracy measures are based on the calls after filtering (that is those calls with a PASS field in the VCF. Parliament2 will still discover calls beyond this filter set, which could be re-validated through other methods to recover more SV events without losing precision.

**Figure 7.**
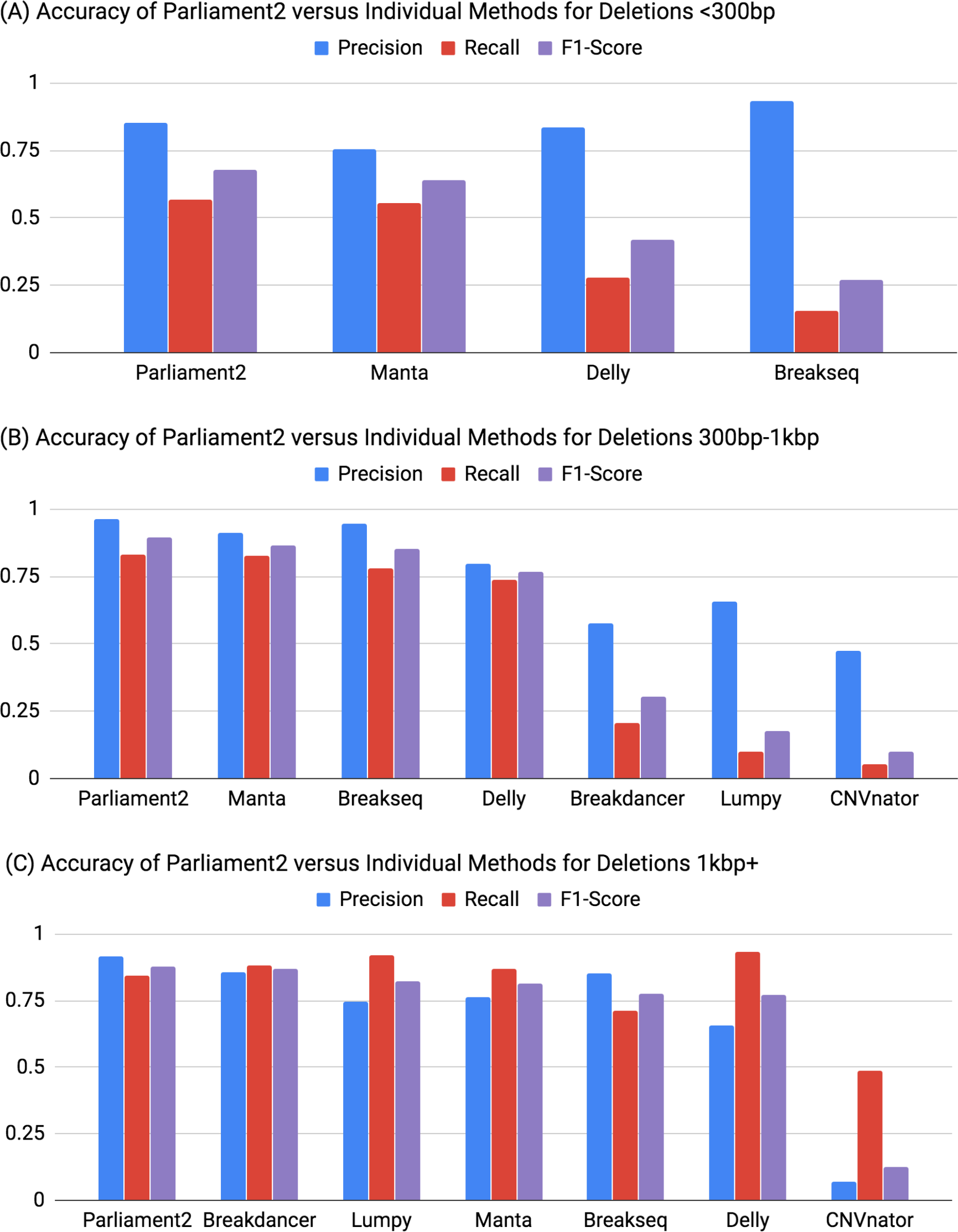
Accuracy of Parliament2 final calls compared to individual tools. The figure above indicates the precision, recall, and F1-score for Parliament2 and the individual methods that compose it, stratified across deletions (A) less than 300 bp, (B) between 300 bp and 1 kbp, and (C) larger than 1 kbp. The order of methods in each graph is sorted such that methods with higher F1-score are located toward the left. The power of individual methods relative to each other can vary substantially between size ranges.

### Generating a Structural Variant Callset on 1000Genomes for GRCh38

To demonstrate the scalability of Parliament2 for large datasets, and to create a resource of value for understanding SVs, we applied Parliament2 to analysis of the low coverage WGS samples from the 1000Genomes project. Structural variants have previously been called on GRCh37 reference [1]. Although the 1000Genomes has been remapped to GRCh38[22], we are not aware of a project-wide set of SV calls from multiple routines using these data.

The computational requirements were modest by comparison to other familiar applications and the entire SV calling was completed in one day of wall clock time, using only 63,720 CPU-hours (about 24 core-hours per sample). For reference, that amount of compute is approximately equivalent to running GATK4 on 220 WGS samples at 35X coverage. This created calls for each of Breakdancer, CNVnator, Delly, Lumpy, and Manta, as well as SVTyped files of each and their union.

In total, there are 88,404 deletion events greater than 50 bp discovered in this set and 30,479 inversion events (Figure 8). There were only 619 insertion events discovered, possibly reflecting that it is more difficult to assemble an insert in low coverage data. The number of calls per sample was generally lower than observed for the high-coverage WGS samples investigated in the prior benchmarks. In addition, for certain samples some of the callers did not generate any output, possibly due to low sequence coverage of the samples.

**Figure 8.**
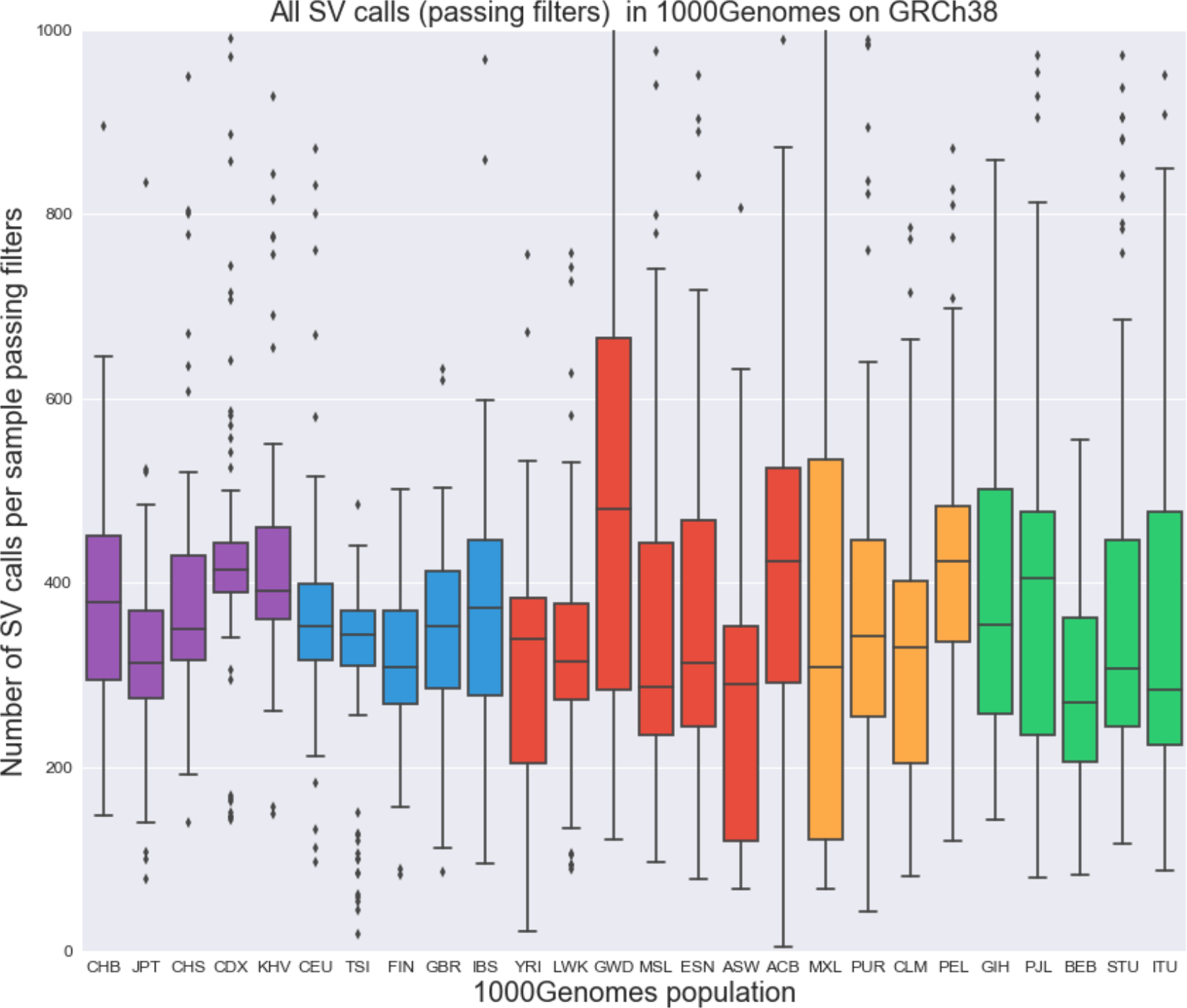
Population distribution of SV calls produced by Parliament2 for the 1000Genomes. The figure above shows the distribution of all SV calls which pass quality filters per sample in the 1000genomes. Populations are colored by their super population code: EAS-purple, EUR-blue, AFR-red, AMR-orange, SAS-green.

These SV calls will be invaluable for understanding the differences between GRCh38 and GRCh37, to provide a resource to understand SVs called on GRCh38 in unknown samples relative to 1000Genomes and to understand how each of these tools interacts with the low coverage data in 1000Genomes.

Download links for the following resources are:

A project-level VCF of all PASS variants in any sample: https://dl.dnanex.us/F/D/J4GVpbx9qjbGG1v3jV00KVg4yQfx8P671PpK2PxP/1000genomes.parliament.pass.vcf.gz

A tar.gz file containing the combined Parliament2 output files: https://dl.dnanex.us/F/D/b879F65ZYBxYqJ47PzKvv32F7Z64fXJ263KF5qPP/parliament2.combined.genotyped.tar.gz

A tar.gz file containing all Breakdancer calls: https://dl.dnanex.us/F/D/X5YpGFF7Xv9GV6Z00J531z6KyK576vqB7FY8G90B/parliament2.breakdancer.tar.gz

A tar.gz file containing all CNVnator calls: https://dl.dnanex.us/F/D/KJkGzq614YkQFp6zXxxbfY78KxJPq0Pjx6p035gp/parliament2.cnvnator.tar.gz

A tar.gz file containing all Delly calls: https://dl.dnanex.us/F/D/pkGjQPkb9yzxbp120PfgJZY2qGXPpvk8FX754KP5/parliament2.delly.tar.gz

A tar.gz file containing all Lumpy calls: https://dl.dnanex.us/F/D/vk227zpkXz43jVvfF2Qg1kvqvGFz6pGFbXZQq43G/parliament2.lumpy.tar.gz

A tar.gz file containing all Manta calls: https://dl.dnanex.us/F/D/81VPqy7gvGYv60zPQqjpXG15KPgk155gf51KVPGy/parliament2.manta.tar.gz

## Discussion

Parliament2 allows structural variant discovery with high recall across a wide size range of types of SVs including duplications, deletions, insertion, inversions and translocations. One of the remaining challenges in SV detection is to simultaneously archive high recall and sensitivity. While SNP calling often achieves this on certain benchmark sets, SVs callers often suffer from either being too sensitive, sacrificing precision or else having a high precision at the expense of sensitivity. Given our newly developed quality scoring we can now leverage the benefits of different approaches/heuristics to ensure a higher quality (precision) and recall. Furthermore, Parliament2 is highly tunable allowing the incorporation of additional methods as well to only run a subset of the methods. As a result, Parliament2 will likely be useful in the distant future when SV methods mature and improve in their prediction.

A unique feature of Parliament is the ability to tune the precision/sensitivity to cope with different scenarios. For example, research discovery has different requirements to clinical applications. The quality of the Parliament2 method also depends on the available DNA sequence coverage per sample. Although the method was developed with an expectation of a minimum of 20x, we illustrated successful processing of low coverage data from 1000Genomes.

The methods are implemented in a computationally efficient mode. This initial call set including genotyped SVs was produced within 24 hours using a tiny fraction of the initial costs for computing the SVs calls in phase3 of the 1000genomes project. This highlights the power of Parliament2 to scale up to high sample sizes.

We compared the performance of each component as well as Parliament2 to the current version (v0.6) of the NIST Giab SVs calls. The comparisons revealed the strengths and weaknesses of the current methods and showed that the combination of methods can provide high accuracy. For Parliament2 we did not precisely overlap all SVs that were identified by GiaB. It is important, however, to note that the GiaB call set was generated with multiple technologies (Pacific Biosciences, 10X, BioNano, Nabsys, and Oxford Nanopore) and approaches (mapping based and de novo assembly based) whereas Parliament2 utilized short reads only.

Overall Parliament2 represents a highly tunable, fast and accurate methods to infer SVs based on short reads. Its dexterity is enhanced by its availability via Docker images and its changeable interface. Parliament2 Is scalable and freely avialable.

With the number of sequenced genomes expanding dramatically with every year, Parliament2 will ensure that a large body of structural variant callsets can be readily and cheaply generated to ensure that our ability to understand structural variation within the datasets grows faster than our ability to generate the data.

## Methods

### Parliament2 Implementation

The code for Parliament2 is in a github repository with an open-source (Apache-2.0) license at the 1.0.7 version (commit 97517b1a22104a3e0a0966a79c3b5556fde8a89d). Execution of Parliament2 occurred by running v1.0.7 of the Parliament2 DNAnexus app (app-FJ8Fj88054JxXFygKvFqQ39j), which is publicly available to run by any user on DNAnexus. This app runs a docker image built directly from the Github repository. Executions of the app with user-provided input for tool combinations specify the parameter flags to that Docker image to include or exclude the desired tools.

### Input WGS Data Used for Timing and Accuracy Benchmarks

Timing statistics and resource utilization was determined by executing the Parliament2 app on a 35X WGS sample for HG002 that was made by random downsample of the 50X PCR-Free HG002 HiSeqX sample generated for the Challenge set of the PrecisionFDA Truth Challenge. This file is uniquely identified on DNAnexus as file-F4JFyx00Fq4YZkB80P7fZ4Fv and can be downloaded: https://dl.dnanex.us/F/D/3YfQGKfZ4q90pQz84jJQG5zQ8j8XY1g8317BVFgX/HG002-NA24385-50x.70_percent.markdup.realigned.bam (note that this file is 73GB before downloading).

### Timing for individual tools and Parliament2 combinations

All timing calculations are run on a c3.4xlarge AWS instance (16-core, 30GB RAM, 320GB disk). To calculate the runtime and resource utilization of individual components, the Parliament2 app was launched with the desired tool or tool combinations. DNAnexus apps write an entry of machine resource (CPU percent, RAM, and disk utilization) every 10 minutes to a job log which also contains the stdout and stderr outputs for job execution. All info log entries of this after the stderr line for program execution up until the SVTyper step (which indicates completion of all jobs) were taken to determine the resource plots over time.

These execution logs can be retrieved without incurring any cost nor needing a DNAnexus account by installing the DNAnexus open-source SDK at https://wiki.dnanexus.com and issuing the command: dx watch -a nAPM5RFEdGeFI1hkW08caWIV3l6oWssW <job-id>, where job-id corresponds to one of the execution ids:

Breakdancer-only:job-FJ6Y8x804KF01f5GP75f95qk.Breakseq-only:job-FJ6Y9J804KF1y5J1589k9g85, CNVnator-only:job-FJ6YG0804KF3G1Z434vGjVGJ, Delly-only:job-FJ6Y9pQ04KF9JzzG5K1Gy266, Lumpy-only:job-FJ6YGZQ04KF6bz4X65k4FB71, Manta-only:job-FJ6YJ3Q04KFPxyp821XKjq5F, Breakdancer&Manta:job-FJ6YJBQ04KF1zpq13BFK44Gx Breakdancer&Manta&CNVnator:job-FJ6YK3j04KF1zpq13BFK44J4, Breakdancer&Manta&CNVnator&Breakseq:job-FJ6YKB004KF1z2K99Z3FZKvG, Breakdancer&Manta&CNVnator&Breakseq&Lumpy:job-FJ6YPBj04KF9VkVx6Z68g6Jv, Breakdancer&Manta&CNVnator&Breakseq&Lumpy&Delly:job-FJ6YPf804KF8JvV76Y6q8G6j

The -a <token> uses a VIEW only persistent token that gives anyone using the DNAnexus SDK the ability to describe jobs, logs, and objects in the project used for this investigation. The input parameters for the job can be retrieved in a similar way with dx describe -a nAPM5RFEdGeFI1hkW08caWIV3l6oWssW <job-id>.

These analyses may be reproduced by relaunching on the same machine type and parameters with **dx run –clone <job-id>**; however, this must be launched in your own project and this will incur expense for the machine.

### Accuracy comparisons

Accuracy comparisons are performed using Truvari[19] with the following execution: truvari.py -b GIAB_DEL0.6.vcf.gz -c <parliament_output> -o <output_directory –passonly –includebed GIAB_0.6.bed –pctsim=0 -r 2000 –giabreport

To determine accuracy for size ranges, -s <lower_size> and -S <upper_size> were used. The deletion truth set was taken by extracting SVTYPE=DEL from the v0.6 truth set. The insertion truth set was taken similarly by extracting SVTYPE=INS.

The Genome in a Bottle data for v0.6 truth set is at: ftp://ftp-trace.ncbi.nlm.nih.gov/giab/ftp/data/AshkenazimTrio/analysis/NIST_SVs_Integration_v0.6/

### Tool Versions

The individual tools which compose Parliament2 run the following versions of each program:

Breakdancer: [v1.4.3]

(https://github.com/genome/breakdancer/releases/tag/v1.4.3)

BreakSeq2:[v2.2]

(http://bioinform.github.io/breakseq2/)

CNVnator:[v0.3.3]

(https://github.com/abyzovlab/CNVnator/commit/de012f2bccfd4e11e84cf685b19fc138115f2d0d)

Delly:[v0.7.2]

(https://github.com/dellytools/delly/releases/tag/v0.7.2)

Lumpy: [v0.2.13]

(https://github.com/arq5x/lumpy-sv/commit/f466f61e02680796192b055e4c084fbb23dcc692)

Manta: [v1.4.0]

(https://anaconda.org/bioconda/manta)

